# Sex dependent effect of amyloidosis on functional network ‘hub’ topology is associated with downregulated neuronal gene signatures in the APP*swe*/PSEN1dE9 double transgenic mouse

**DOI:** 10.1101/2024.05.13.593982

**Authors:** Zachary D. Simon, Karen N. McFarland, Todd E. Golde, Paramita Chakrabarty, Marcelo Febo

## Abstract

**Background:** Extracellular amyloid-β (Aβ) impairs brain-wide functional connectivity, although mechanisms linking Aβ to broader functional network connectivity remain elusive.

**Objective:** Here, we evaluated the effect of Aβ on fear memory and functional connectome measures in mice.

**Methods:** Middle-aged (9-11 months of age) double transgenic APP-PS1 mice and age and sex-matched controls were evaluated on a fear conditioning protocol and then imaged at 11.1 Tesla. Brains were harvested and processed for analysis of Aβ plaques and Iba1 immunolabeling in cortex, hippocampus, and basolateral amygdala. Additional RNA sequencing data from separate age, strain, and sex matched mice were analyzed for differentially expressed genes (DEGs) and weighted gene co-expression networks.

**Results:** In both male and female mice, we observed increased functional connectivity in a dorsal striatal/amygdala network due to Aβ. Increased functional connectivity within this network was matched by increases in AβPP gene expression, Aβ and Iba1 immunolabeling, and an upregulated cluster of DEGs involved in the immune response. Conversely, the network measure representing node ‘hubness’, eigenvector centrality, was increased in prefrontal cortical brain regions, but only in female APP-PS1 mice. This female specific-effect of amyloid was associated with downregulation of a cluster of DEGs involved in cortical and striatal GABA transmission, anxiogenic responses, and motor activity, in female APP-PS1 mice, but not males.

**Conclusions:** Our results contribute to a growing literature linking between Aβ, immune activation and functional network connectivity. Furthermore, they reveal effects of Aβ on gene expression patterns in female mice that may contribute to amyloidosis-induced dysregulation of non-cognitive circuitry.

## Introduction

Early onset Alzheimer’s disease (AD) has an estimated prevalence of 1-5%, with the remaining 95% of all cases representing late onset sporadic forms ^1^. Approximately 200,000 individuals in the US suffer symptoms as early as 30-40 years of age, compared to non-familial AD which emerges at or after 65 ^2^. Studies of isolated familial cases have led to the discovery of several heritable amyloid-β protein precursor (AβPP) mutations and over 200 highly penetrant mutations in presenilin1 (PSEN1) that are causative for a percentage of early onset cases ^3, 4^. Familial AD involves mutation-specified amino acid substitutions in N- or C-terminal domain residues, or in Aβ regions of AβPP that causes aberrant AβPP processing leading to increased Aβ levels, aggregation and extracellular Aβ plaque formation^3^. The Swedish AβPP (AβPP*swe*) mutations, for example, produces amino acid substitutions (K670N, M671L) near the beta-secretase cleavage site of AβPP, leading to increased production of a 42 amino acid long Aβ peptide (Aβ42) ^4^. Deletion of exon-9 in the PSEN1 gene sequence impairs gamma-secretase activity, which results in formation of aggregation prone Aβ42 ^4^. Aβ is thought to mediate functional deficits in neurotransmission, including changes adversely impacting the activity of both excitatory and inhibitory neurons ^5, 6^. Understanding these and other Aβ-linked functional changes, at the genetic, neuronal and neural circuit scales, is fundamental to uncovering targetable early pathology and neural signatures driving behavioral and cognitive changes in early onset and sporadic forms of AD.

Many reports on changes in excitatory and inhibitory neurons and synapses in AD brains are extrapolated from postmortem assessments ^6–9^. These neuronal changes, and their broader effects on network activity in AD, has been further explored in mice harboring human familial AβPP mutations ^10, 11^ and resting state functional magnetic resonance imaging studies (fMRI) ^12^. Functional neuroimaging studies of early onset AD have reported widespread functional connectivity declines across multiple well-established networks ^13^, particularly involving default mode network (DMN) regions ^14^. This includes lower connectome measures of network strength, efficiency and clustering in frontoinsular, parietal, temporal and occipital cortices ^15^, and reduced eigenvector centrality in bilateral medial temporal gyrus, right posterior hippocampal gyrus, and right postcentral gyrus ^16^. Eigenvector centrality has emerged as an important marker of functional connectomes measured in normal aging, mild cognitive impairment, and AD^16–18^. This graph theory measure correlates with cerebrospinal AB levels, tau, and is linked to cognitive performance in AD and in normal aging^17, 19–22^, although this has not been observed in other studies^23^. Indeed, there is some disagreement between studies in terms of the specific regions affected and extent of reductions in functional connectivity and connectome measures. Pini *et al* reported a greater reduction in hippocampal connectivity to a group of cortical memory related regions in late onset versus early onset AD ^24^. Adriaanse *et al* observed broad functional connectivity declines in early onset AD compared to late onset, the latter mostly had reductions in the DMN compared to controls ^13^. Yet, others have reported only DMN reductions relative to controls in early onset AD ^14^. All the above studies use similar aged subjects, around the age of sixty and matched control groups. Although pathology of AD is highly characterized by amyloidogenic mechanisms related to AβPP and PSEN mutations, there are other disease contributing genes differentially expressed in this population ^25, 26^. Moreover, there are significant sex differences in the genetic architecture conferring resilience to AD, particularly cognitive resilience ^27^. How this translates to functional brain changes in males versus females in response to AD pathologies remains an area of active research. Thus, the oligogenic contributions to AD, and important biological factors such as sex, warrant consideration in experiments to examine mechanistic links between Aβ and brain functional connectivity.

Functional neuroimaging studies in transgenic rodents harboring AβPP, PSEN1 and other AD causative or moderate-to-high risk mutations, selectively or in combination, can help clarify specific gene-pathology-brain functional connectivity interactions. Several animal fMRI studies have begun to address Aβ-functional connectivity interactions, and these shown results consistent with some human studies of early onset AD ^28–31^. Mouse fMRI studies have revealed nuanced interactions between Aβ aggregation and functional connectivity outcomes ^30^, which calls for attention to how AβPP is processed and how it interacts with other AD genes. Here we report new findings in the APP-PS1 mouse ^32^ to expand on the existing literature on the relationship between Aβ and functional connectome changes. Further, we report a potentially important link between female-specific effects of Aβ on functional connectome node eigenvector centrality and brain gene expression. The differentially expressed genes specifically affected by Aβ in females include genes in involved in striatal inhibitory gamma amino butyric acid (GABA) transmission.

## Methods

### Animal studies

Mice were housed in age- and sex-matched groups of 3-4 in a temperature- and humidity-controlled room, inside conventional air filtered cages (dimensions: 29 x 18 x 13 cm) with food and water available *ad libitum* (vivarium lights on from 07:00-19:00 hours). Transgenic (TG) cohorts were generated in house by breeding heterozygous male mice bearing amyloid-β protein precursor (AβPP) with the Swedish familial mutation and exon 9 deficient presenilin-1 (PSEN1) and hybrid C57BL6/J/6xC3H F1 females (Envigo)^32^. APP-PS1 mice develop gliosis and plaque accumulation starting at 6 months of age and long-term cognitive impairment as early as 2 months of age^33^. Control mice were littermate non-transgenic (NTG) mice, and all groups contained male and female mice. Experiments started when mice were 10-11 months of age, which corresponds to high amyloidosis, gliosis, with previously reported impairments in cognitive function and other non-cognitive behaviors ^34^. The behavior and imaging experiment group sizes were 13 NTG controls (including 4 female NTG) and 15 TG mice (including 7 female TG). The tissue histology group sizes differed between regions based on slice quality. Groups were 8-9 NTG controls (including 3-4 female NTG) and 11-13 TG mice (including female 5-7 TG). All procedures received prior approval from the Institutional Animal Care and Use Committee of the University of Florida and follow all applicable NIH guidelines.

### Fear conditioning

Fear conditioning was performed two weeks before fMRI studies to obtain a readout related to memory function and emotion. In addition to offering a screening for aspects of these behavioral dimensions, it offered a short length of training and testing compared to other commonly used behavioral assays, such as Y-maze spatial alternation and Morris Water Maze tests. The fear conditioning protocol was modified from^35^ and previously described in^36^. Mice were monitored in a test cage for freezing behavior (e.g., transient bouts of immobility) during initial exposure to an auditory stimulus paired with a mild electric current and then monitored once again at 24 and 48 hours in the presence of the auditory cue. The percentage of freezing time in response to the auditory cue was used as a measure of conditioned fear.

The fear conditioning cages were housed in sound attenuation chambers. The chambers had computer-controlled lighting, white noise generator, a fan and control units. These included audio speaker controls for tone generation, shock controllers for the internal cage floors, and infrared and white lighting controllers. The internal operant cage was made of translucent Plexiglas with a steel frame and steel shock grid floor connected to a computer controlled current stimulator (Med-Associates, Inc. St. Albans, VT). Activity within the fear conditioning cages was captured by a monochrome video camera. All cage accessories were controlled by PC running Video Freeze Software^TM^. Camera lens brightness, gain, and shutter settings were verified and adjusted before collecting data and were kept constant across all mice. For each session, the detected motion-sensitive signals were calibrated with the mouse inside the cage and NIR lights on. The estimated motion index was set to the same threshold level across all mice, prior to data exporting or analyses. Videos were acquired at 30 frames per second (fps) with a minimum freeze duration of 30 frames.

On day 1 (training), mice were exposed to 4 consecutive presentations of a tone (20 second duration, 90 dB tone, 5kHz, 50ms risetime; 190-second inter-tone interval), each of which ended with the presentation of a brief 1-second 0.9mA current pulse to the entire floor of the cage. The same tone presentation protocol was carried out in subsequent tests at 24- and 48-hours but in the absence of the electric stimulus. In addition, the cage environment was modified on the 48-hour test by placing a plastic smooth white floor and diagonal walls, which covered the grid floor, light, and speaker. Thus, trials on the training session (day 1) measure the degree to which mice learn to associate the sound (conditioned stimulus) with the electric current (unconditioned stimulus) through an increase in freezing behavior (unconditioned response). The level of freezing behavior (conditioned response) at 24 hours indexes the degree of recall or memory of the CS-UCS association. Modifying the cage conditions at the 48-hour test enabled measuring the contribution of context to the conditioned response. Across all days of testing, freezing behavior was assessed by measuring immobility time within a 60-second window during the presentation of the auditory tone (which includes the 20 second tone epoch and an additional 40 seconds after the end of the tone). For each of the epochs corresponding to the 4 tone-shock pairings, a component comprising a 20 second pre-shock tone interval and a 40 second post-shock interval were used to quantify percent component freezing (per day there were four 60 second components summed to provide total percent freezing per day). Motion-index was used as a surrogate measure of locomotor activity.

### Magnetic resonance imaging Instrumentation

Images were collected on a magnetic resonance spectrometer tuned to 470 MHz proton resonant frequency (Magnex Scientific 11.1 Tesla/40 cm horizontal magnet). The MRI system was equipped with Resonance Research Inc. spatial encoding gradients (RRI BFG-240/120-S6, maximum gradient strength of 1000 mT/m at 325 Amps and a 200 µs risetime) and controlled by a Bruker AV3 HD console running Paravision 6.0.1 (Billerica, MA). A quadrature transceiver surface radiofrequency (RF) was used for B1 signal generation and detection (RF engineering lab, Advanced Magnetic Resonance Imaging and Spectroscopy Facility, Gainesville, FL).

### Mouse imaging setup

Mice were scanned sedated under a continuous paranasal flow of 0.25 % isoflurane gas (delivered at 0.5 L/min mixed with medical grade air containing 70% nitrogen and 30% oxygen, Airgas, Inc.) and a continuous subcutaneous infusion of dexmedetomidine. Prior to setup, mice were anesthetized under 2% isoflurane and administered an intraperitoneal (i.p.) injection of 0.1mg/kg dexmedetomidine (at a 1ml/kg volume). Isoflurane was reduced to 0.25% throughout the remaining imaging session. An infusion line was set up for subcutaneous delivery of dexmedetomidine over the course of scanning (0.1mg/kg/ml at an infusion rate of 25 µl/hour using a PHD-Ultra microinfusion pump, Harvard Apparatus, Holliston, MA). Functional MRI scans were collected at least 50 minutes after the i.p. injection. Respiratory rates were monitored continuously, and temperature was controlled remotely using a warm water recirculation system (SA Instruments, Inc., New York).

### Functional MRI acquisition

We acquired a T2-weighted anatomical and an fMRI scan per mouse. The T2-weighted Rapid Acquisition with Relaxation Enhancement (RARE) sequence was acquired with the following parameters: echo time (TE) = 41 ms, repetition time (TR) = 4 seconds, RARE factor = 16, number of averages = 12, field of view (FOV) of 15 mm x 15 mm and 0.8 mm thick slice, and a data matrix of 256 x 256 (0.06 µm^2^ in plane) and 12 interleaved ascending coronal (axial) slices covering the entire brain from the rostral-most extent of the anterior prefrontal cortical surface, caudally towards the upper brainstem and cerebellum. Functional images were collected using a single-shot spin echo planar imaging (EPI) sequence with the following parameters: TE = 16 ms, TR = 1.5 seconds, 600 repetitions, FOV = 15 x 15 mm and 0.9 mm thick slice, and a data matrix of 64 x 48 (0.23 x 0.31 µm in plane) with 14 interleaved ascending coronal slices in the same position as the anatomical scan. Ten dummy EPI scans were run prior to acquiring data under steady state conditions. Respiratory rates, isoflurane and dexmedetomidine delivery, temperature, lighting, and room conditions were kept constant across subjects.

### Image pre-processing

The image processing workflow followed previously published research ^36–40^. Resting state processing was carried out using Analysis of Functional NeuroImages (AFNI) ^41^, FMRIB Software Library (FSL) ^42^, and Advanced Normalization Tools (ANTs) ^43^. Binary masks for anatomical and functional scans were created using ITKSNAP ^44^. The brain binary masks were used for brain extraction prior to registration steps. Times series spikes were removed (3dDespike, AFNI), image repetition frames aligned to the first time series volume (3dvolreg, AFNI), and detrended (high pass filter <0.009 Hz using AFNI 3dTproject). Independent component analysis (ICA) using Multivariate Exploratory Optimized Decomposition into Independent Components (FSL MELODIC version 3.0) was used to assess structured ‘noise’ or artefact components in each subject scan, in their native space. Most, if not all ICA components in this first stage contain artefact signal voxels along brain edges, in ventricles, and large vessel regions. These components were suppressed using a soft (‘non-aggressive’) regression approach, as implemented in FMRIB Software Library (FSL 6.0.5) using fsl_regfilt ^42^. A low-pass filter (>0.12Hz) and spatial smoothing (0.4mm FWHM) were next applied to the fMRI scans prior to registration steps. Post-regression ICA was carried out to verify removal of artefact components and preliminary assessment of putative ICA networks in individual scans.

### Atlas registration and resting state signal extraction

Anatomical scans were cropped and bias-field corrected (N4BiasFieldCorrection, ANTs). Functional scans were cropped, and a temporal mean image was registered to the anatomical. Preprocessed anatomical and fMRI scans were aligned to a parcellated mouse common coordinate framework (version 3, or CCFv3) template ^45^. Bilateral region of interest (ROI)-based nodes (148 total) were created with the guidance of the annotated CCFv3 parcellation and using tools in ITKSNAP and FSL, similar to our previous work in rats ^46, 47^. Large brain regions, such as hippocampus, motor, somatosensory, and visual cortices, were assigned multiple nodal masks. These are distinguished based on atlas coordinates and a numerical identifier at the end of the regional code (e.g., hippocampus contains nodes HPC1-HPC5). Anatomical images were linearly registered to the mouse template using FSL linear registration tool (FLIRT), using a correlation ratio search cost, full 180-degree search terms, 12 degrees of freedom and trilinear interpolation. The linear registration output was then nonlinearly warped to template space using ANTs (antsIntroduction.sh script). Anatomical-to-atlas linear and nonlinear transformation matrices were applied to fMRI scans at a later stage. Spontaneous signals, each with 600 data points, were extracted from 148 ROIs and used in cross-correlations and in calculations of Pearson r coefficients for every pairwise combinations of ROIs (using *corrcoef* function in MATLAB). The resulting number of pairwise correlations was 10,730 per subject (after removing 148 self-correlations). Correlation coefficients were Fisher’s transform to ensure normality prior to statistical analyses. The center voxel coordinates for the ICA-based nodes normalized to the CCFv3 were used in 3D network visualizations using BrainNet viewer ^48^.

### Network analysis

Weighted undirected matrices were analyzed using Brain Connectivity Toolbox ^49^ in MATLAB (Mathworks, Natick, MA). Global graph metrics were calculated for edge density thresholds ranging from 2-40%. Global network measures for this density range were converted to area under the curve (AUC) values prior to statistical assessments. Node-specific network measures were converted to AUC values per node. We assessed several graph measures of network integration and communication efficiency in APP-PS1 and control mice. This included node strength (sum of edge weights/node) and degree (sum of edges/node) and global measures, such as transitivity (related to CC; number of triad groups normalized by all possible triad nodes in a network), and characteristic path length (CPL; the average edges or edge weights between node pairs). For local node efficiency, a length matrix was used to calculate inverse values in vicinity of a node, with added weights used to emphasize the highest efficiency node paths ^49, 50^. To corroborate results relative to random networks, all network measures were calculated on original and randomized versions of the functional connectivity matrices. Positive and negative edges were randomized by ∼5 binary swaps with edge weights re-sorted at each step. A probabilistic approach for community detection was used to calculate a modularity statistic (Q), which indexes the rate of intra-group connections versus connections due to chance ^51^. The procedure starts with a random grouping of nodes and iteratively moving nodes into groups which maximize the value of Q. The final number of modules and node assignments to each group (e.g., community affiliation assignments) was taken as the median of 100 iterations of the modularity maximization procedure ^46^. We analyzed the tendency of assortative vs dissortative mixing of nodes ^52^Δ. The assortativity index is a correlation coefficient comparing node strength values between pairs of edge-connected nodes. Positive r values indicate connectivity between pairs of nodes with similar strengths (e.g., high strength nodes pairs with high and low with low), while negative r values indicate cross-pairings between low and high strength nodes. We also analyzed nodal eigenvector centrality and betweenness centrality, as previously reported^46^.

### Immunohistochemical labeling of amyloid plaques and microglial cells

Mouse brains were harvested and processed for immunohistochemistry. Formalin fixed brain hemisections were paraffin embedded and sectioned sagittally using microtome at the following Bregma locations: lateral 0.12mm, 1.56mm, and 3.25mm. Consecutive 4-μm were collected at each level for staining. The primary antibodies were Iba1 (1:1000, Wako Catalog # 019-19741) and the Aβ1-16 antibody (33.1.1) for staining Aβ plaques (1:13,000; provided by the Chakrabarty laboratory, Center for Translational Research on Neurodegenerative Diseases, University of Florida; kind gift of Todd E Golde, Emory University). Whole slides were scanned on a ScanScope XT Scanner (Leica Biosystems). Images were analyzed with tuned positivity algorithms for each respective stain using the Aperio ImageScope software. Regions were hand-drawn on each image using the Paxinos and Franklin’s Mouse Brain Atlas in Stereotaxic Coordinates 4^th^ ed. as a guide. The images had the following regions segmented out: prefrontal cortex, sensorimotor cortex, and hippocampus (at approx. Bregma coordinates lateral 1.76), and basolateral amygdala (at approx. Bregma coordinates lateral 2.88). The positivity threshold (which highlights the dark-stained tissue beyond a certain threshold) was tuned to highlight the staining of each respective stain, but not the surrounding tissue. Tuning was done on a single image for each stain to obtain the threshold which would produce most signal from only the stained tissue and not surrounding unstained tissue. This threshold for each stain was kept the same for all regions and all subjects throughout the analysis. The obtained positivity values were the percentage of pixels positive for staining at the tuned threshold, within the given drawn region. Each animal has a single positivity value from the specified region drawn on the image of the stained slide containing that region. 2-way ANOVA with main effects and interaction of sex and genotype was used to determine p-values for differences between groups within each specified brain region.

### Normalized gene counts and analysis of differentially expressed genes (DEGs)

Forebrain RNA sequence data of the APP/PS1 mice were obtained from the AD Knowledge Portal synapse.org repository (Project SynID: syn2580853, title: “The Tau and AβPP Mouse Model (TAUAPPms) Study”). The dataset included 9-12 month NTG (n=15, including 7 NTG females) and 12 month TG mice (n=11, including 5 TG females). BAM files were downloaded from Synapse.org as described (doi: https://doi.org/10.7303/syn2580853). Rsamtools and GenomicAlignments R packages ^53, 54^ were used to generate gene counts from the BAM files and were filtered for gene counts to remove genes with low counts (<10 genes). Subsequent analyses used normalized gene counts expressed as fragments per kilobase of transcript per million mapped reads (FPKM values). Analyses of DEGs was carried out using DESeq2 using default setting and a general linear model fit per gene ^55^. Wald tests were used to compare TG vs NTG groups and multiple comparison adjustments of resulting p-values were done with the Benjamini-Hochberg false discovery rate (FDR) correction method. DEGs were defined as an absolute log2 fold change > 1 and an adjusted p value < 0.05 (visually evaluated as volcano plots). Gene annotation symbols were used to group genes according to their known expression in mouse brain cell types ^56^. Within these cell type classifications, the geometric mean of the normalized gene counts for the genes comprising the cell types was calculated and analyzed between experimental groups. The cell type classifications included astrocytes, neurons, oligodendrocyte precursors, newly formed oligodendrocytes, myelinating oligodendrocytes, endothelial cells, and microglia/macrophage cells ^56^.

### Weighted Gene Co-Expression Network Analysis (WGCNA)

The WGCNA R package was used to construct gene correlation networks from the normalized expression data after filtering and removing genes with zero variance ^57, 58^. Plots of scale-free topology fitting and mean connectivity, both as a function of soft-threshold power values ranging from 1-20, were evaluated to select a gene network threshold using the “pickSoftThreshold” function. Networks were constructed all male and all female samples separately. Adjacency matrices for a signed hybrid network were constructed using expression data and the chosen power threshold. The adjacency matrices were converted to topological overlap maps (TOMs) to be used for module network graphs. Hierarchical clustering was used to identify modules with the cutreeDynamic function. A deepSplit setting of two with a minimum module size of 30 was used for all analyses. Dendrograms were evaluated and compared by identifiable color-coded module statistics. Modules containing highly co-expressed genes were merged and co-expression similarity of entire modules were quantified by first calculating eigengenes and then clustering these based on their Pearson correlations. A cluster height cut was 0.25 (correlations of 0.75) was applied to merge similar modules. To quantify module-trait associations we recalculated module eigengene correlations and p values and displayed these as heat maps. Gene-trait and gene-module relationships were analyzed based on gene significance and module membership statistics. Module hub genes were determined with the ‘chooseTopHubInEachModule’ function.

For module preservation testing, female data were used as a reference and male data tested against the reference for low (z < 2), moderate (z>2, z<10), high module preservation (z > 10). We used sex as a comparison trait since preliminary assessments indicated sex differences in DEGs and cell type compositions. Statistics were run as multiExpr data for signed hybrid networks with 200 permutations and pariwise comparisons are made between common genes. Intramodule degree for eigengenes (kME) is used as a module membership statistic plotted against z score converted summary statistics determining significance. All statistical comparisons for preservation p values were Bonferroni-corrected.

### Statistical analysis

Statistical analyses and data visualization were carried out using tools in MATLAB, GraphPad Prism (version 10.2) or ggplot2 in the R statistics package. Unless otherwise stated, statistical analyses used either a two-way full factorial analysis of variance (ANOVA) (genotype x sex, critical α<0.05) or Mann-Whitney tests. Post hoc tests used Tukey’s honest significant difference (HSD) procedure. Where appropriate, FDR correction (q ≤ 0.05) was used. Further analysis of fMRI scans was carried out using probabilistic ICA ^59^ as implemented in MELODIC Version 3.15, part of FSL ^59, 60^ and as previously described for mouse fMRI scans ^36^. The resulting components were overlaid on the CCFv3 mouse atlas template and classified according to the peak z statistic anatomical location. A series of two-stage multiple linear regressions were used to back-project group ICA components to subject-specific time courses (using spatial regressions) and spatial components (using temporal regressions). The subject-specific spatial component maps were then used in statistical comparisons for each component in the FSL randomise tool. The statistical design matrix was generated using the FSL Glm tool. We used a one-way ANOVA general linear model design. Randomization tests were carried out with 500 permutations (corrected p-level for significance is 0.05 and statistical thresholding of maps by threshold free cluster enhancement).

## Results

### Female APP-PS1 mice have a higher amount of amyloid plaques in basolateral amygdala and sensorimotor cortex than males

Histological analyses confirmed Aβ plaque presence across hippocampal, sensorimotor, prefrontal and basolateral amygdala regions in TG mice (Figure 1a and 1c). Additional analyses indicated that female TG mice had a greater amount of Aβ plaques in basolateral amygdala region (Mann-Whitney p = 0.0087) and in neocortical areas corresponding to somatomotor cortex (p = 0.035) (Figure 1b). The hippocampus had a similar trend (p =0.051), and prefrontal cortex was not significantly different between male and female mice.

**Figure 1.**
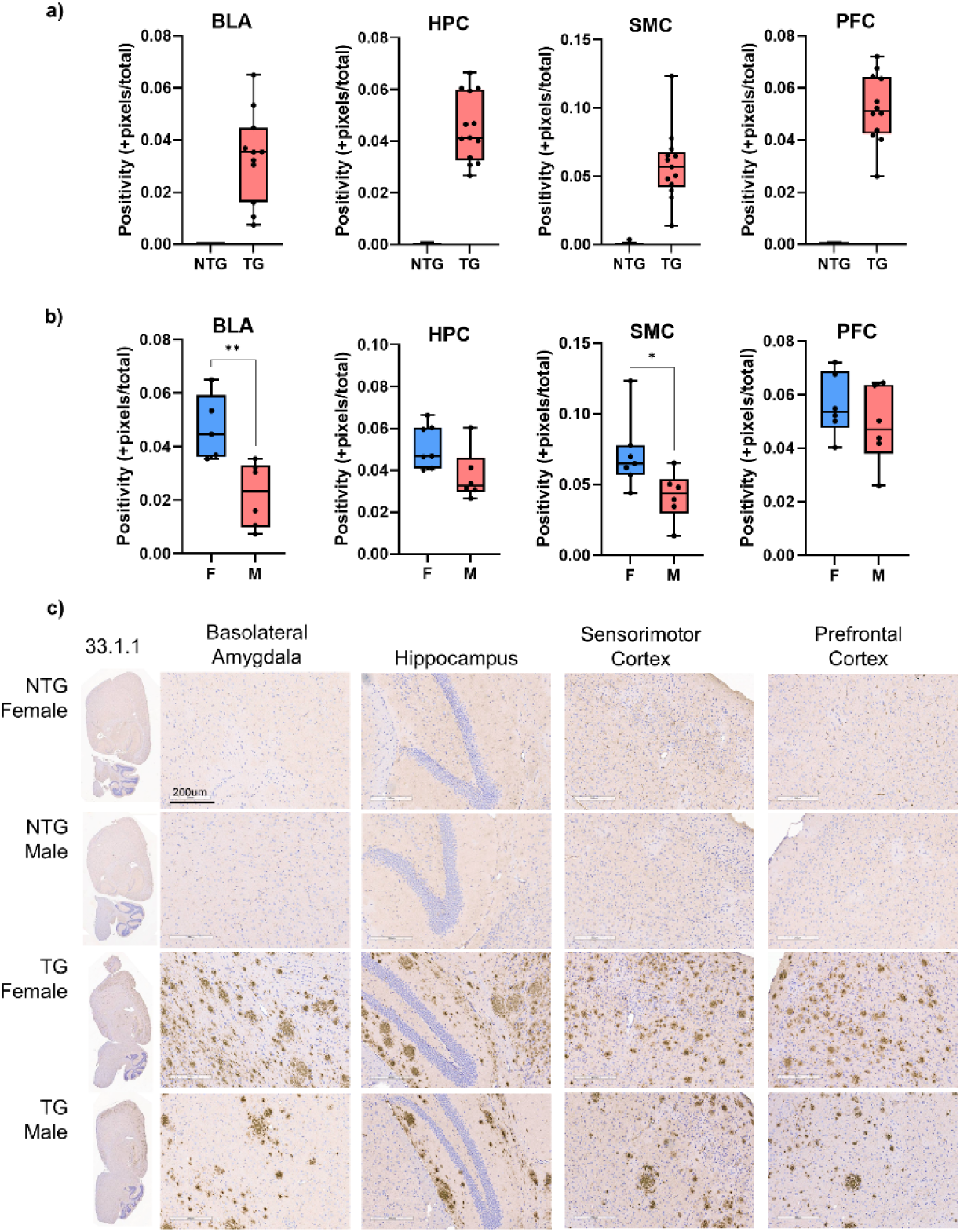
Female APP-PS1 mice have a greater number of amyloid plaques in basolateral amygdala and somatomotor neocortical areas compared to male APP-PS1 mice. a) Confirmation of amyloid plaque presence in TG mice using Aβ1-16 antibody 33.1.1. b) Comparison of amyloid plaque burden between male (M) and female (F) TG mice. c) Representative histological sections from female and male NTG and TG mouse basolateral amygdala (BLA), prefrontal cortex (PFC), somatomotor cortex (SMC), and hippocampus (HPC). Data in a-b are box-whisker showing median, minimum-maximum values and individual data points. Asterisks are *p < 0.05, **p < 0.01.

### Amyloidosis increased functional connectivity within a posterior striatal/amygdala network

ICA identified 20 components that included previously reported networks ^36, 61–63^. These are summarized in Supplementary Figure 1. The components had peak z-statistic voxels in anterior and posterior cingulate, ventral and dorsal striatal, motor, somatosensory, visual cortices, thalamic, midbrain, and cerebellar regions. Statistical analysis using a two factor ANOVA (genotype x sex) indicated no significant main effects or interactions in established ICA networks. We analyzed an unclassified mixed network containing peak z score voxels in the left posterior striatum/basolateral amygdala (Figure 2a). This network included connectivity with cerebellar, prelimbic, motor, posterior cingulate, ventral hippocampus, entorhinal cortex regions. Two-way ANOVA revealed a main effect of amyloidosis, with TG mice having greater functional connectivity across these and other regions compared to NTG mice. No main effect of sex and sex x genotype interactions were observed for this mixed network (Figure 2b). Figure 2b summarizes the brain areas with greater functional connectivity in TG vs NTG mice. These included subregions of the motor cortex, nucleus accumbens, insula, midbrain, piriform cortex, entorhinal cortex and reticular nuclei (p < 0.05, threshold free cluster enhanced corrected).

**Figure 2.**
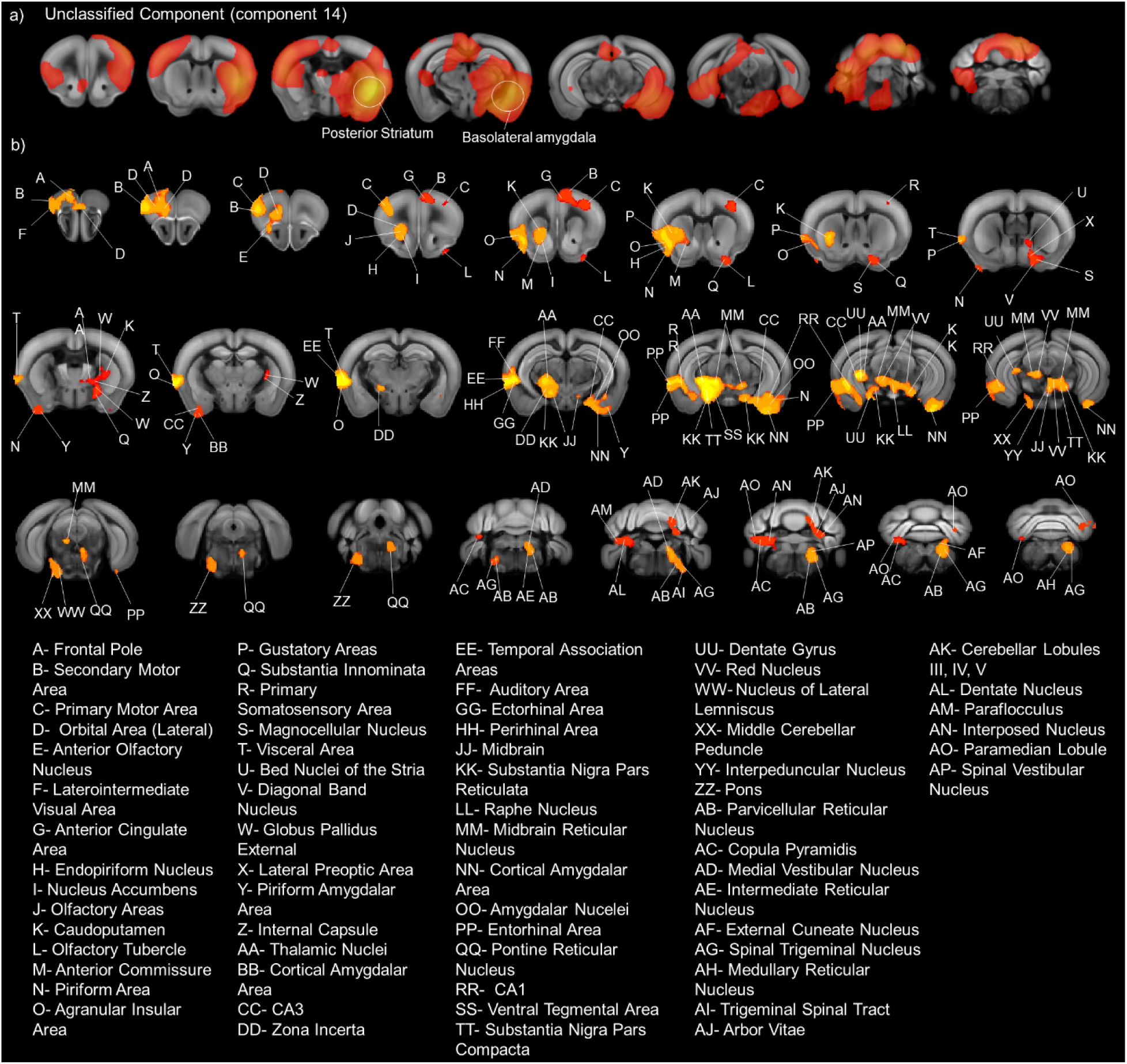
Amyloidosis increased functional connectivity within an unclassified component that included nodes in posterior dorsal striatum and basolateral amygdala in both male and female mice. a) Undistinct component measured in a cohort of 28 NTG and TG fMRI scans. Higher intensity/higher functional connectivity voxels in yellow are encircled and indicated by the corresponding brain region. b) Statistical results of full factorial ANOVA assessing genotype x sex interactions and main effects (p < 0.05, corrected). Letter codes indicate the brain area according to the Allen Mouse Brain Common Coordinate Reference atlas (CCFv3). Brain area names are matched to letter codes at the bottom of the figure. Significant voxels (main effect genotype) are highlighted in yellow-orange hue indicating TG > NTG.

### Amyloidosis is associated with higher eigenvector centrality scores in prefrontal and striatal subregions in female mice

Global network measures did not vary significantly between NTG and TG groups, regardless of sex (Supplementary Figure 2a). Figure 3a shows nodal eigenvector centrality scores calculated at 10% edge density. Eigenvector centrality scores at 10% edge density in several prefrontal and striatal nodes differed as a function of either sex or genotype (in the case of female mice) (Figure 3a). A similar difference was not observed in male mice, which had eigenvector centrality scores similar to TG female mice in these brain areas. TG females had higher eigenvector scores compared to NTG females in a single node in the left prelimbic cortex and two in the right prefrontal cortex, left lateral and ventral nodes in the orbital cortex, a node in the dorsomedial division of the left striatum (group[sex/genotype] x node interaction, F_441,3528_=1.2, p=0.005; multiple between-group comparison t-tests per node with FDR adjusted p values at q < 0.05) (Figure 3a). NTG males had higher eigenvector scores compared to NTG females in the same nodes and in left dorsal agranular insular cortex, right anterior cingulate cortex, right infralimbic cortex, right lateral orbital cortex, right secondary motor cortex, and right dorsomedial striatum (FDR adjusted p < 0.05 for comparisons across all nodes). When eigenvector scores were analyzed as AUC values per node, we also observed similar regions with lower eigenvector centrality scores in NTG compared to TG females and NTG females compared to NTG males (FDR adjusted p < 0.05 for comparison across all nodes). Node eigenvector centrality scores in prelimbic cortex, left lateral orbital cortex, left hippocampus, left dorsal agranular insula, left dorsolateral striatum were greater in TG compared to NTG female mice (Figure 3b). Eigenvector centrality in left and right prelimbic cortex, left hippocampus, right anterior cingulate cortex, and right orbital cortex were greater in NTG males than NTG female mice (Figure 3b). It should be noted that both left and right hemispheres of these brain regions had similar differences between male and female NTG and female NTG compared to female TG (uncorrected p values < 0.05). However, only the mentioned brain regions were significant after FDR adjustment of p values. The summary 3D functional connectome maps in Figure 3c illustrate the average AUC eigenvector score brain distribution in each group.

**Figure 3.**
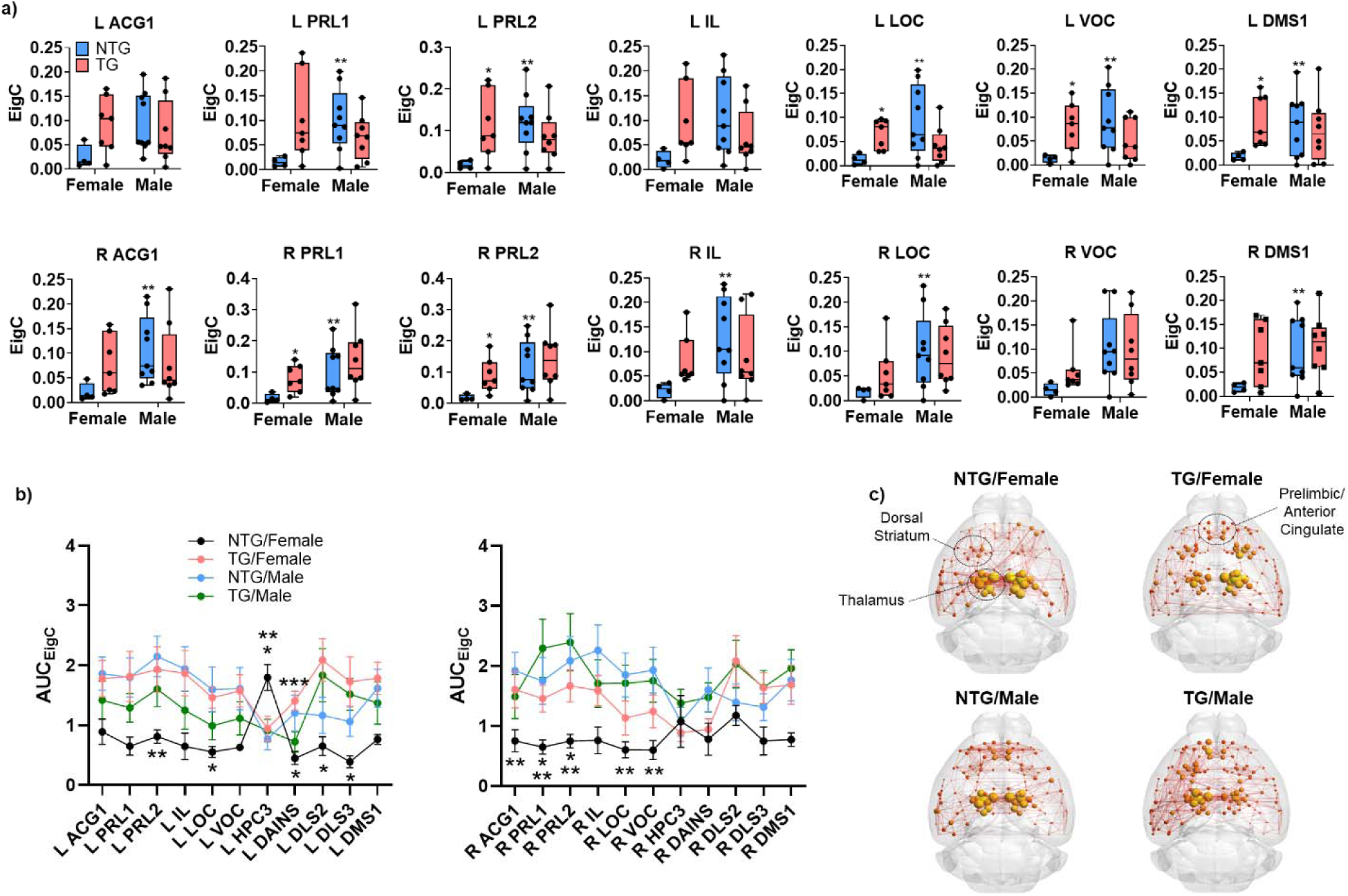
Sex specific effects of amyloid on functional network ‘hub’ topology. a) Eigenvector centrality scores for several prefrontal cortical and striatal nodes calculated for a 10% network density. Data in box-whisker plot are median with minimum and maximum values and show individual data points. Top row plots show ROI data for left (L) hemisphere and bottom row plot the right (R) hemisphere. b) Area under the curve (AUC) values calculated for eigenvector centrality across multiple edge densities (2-40%) per ROI. Data in plot is presented as mean standard ± error. Left plot shows ROI data for left (L) hemisphere and right plot the right (R) hemisphere. c) Connectome maps represent mean AUC values for eigenvector centrality per node and edge strength (Pearson correlations) as spherical nodes and lines connecting nodes. Node eigenvector centrality is reflected in sphere size and color intensity. Edges are thresholded at 6% density for visual purposes. Asterisks indicate significantly difference between NTG vs TG females* and NTG females vs males**. Abbreviations: ACG1, anterior cingulate gyrus; PRL, prelimbic cortex; IL, infralimbic cortex; LOC, lateral orbital cortex; VOC, ventral orbital cortex; DMS, dorsomedial striatum.

### Within-sex effects of amyloidosis on conditioned fear learning and recall

No differences in fear learning were observed between groups (Supplementary Figure 3). Overall behavioral data was analyzed using 2-way ANOVA of main effects and interactions of genotype and trial, with Tukey post-hoc analysis for multiple comparisons. Both TG and NTG mice increased the percentage time freezing over the course of shock-tone exposure trials on the first day of testing (Supplementary Figure 3a). Comparison across days also failed to show any differences between NTG and TG mice on any day (Supplementary Figure 3b). No differences in motor activity were observed on any of the three test days (Supplementary Figure 3c). Within-subject analysis of day one of fear conditioning did reveal differences between groups, including sex differences. First, as expected, we observed an effect of trial on day 1 of testing (main effect of trial, F_3,72_ = 8.8, p < 0.0001). However, Tukey’s multiple comparison test indicated that only male mice developed fear conditioning when comparing trials 3 and 4 to the first shock-cue pairing (NTG males: p = 0.0047 trial 1 vs. 3 and p = 0.0017 trial 1 vs. 4; TG males: p = 0.03 trial 1 vs. 3 and p = 0.02 trial 1 vs. 4). Percent time freezing in trials 3 and 4 compared to the first trial were not significant in NTG and TG female mice (Supplementary Figure 3a). Within-subject analyses also revealed significant differences in fear conditioning over the 3 test days (main effect of day, F_2,48_ = 17.4, p < 0.0001; Supplementary Figure 3b). Percent time freezing was maintained across days in NTG males. Male TG mice showed an increase in percent freezing time on days 2 and 3 relative to day 1 (p = 0.0035 and p = 0.01, respectively). Female NTG and TG mice both showed increases in percent time freezing on day 2 compared to day 1 (female NTG, p = 0.0027 for day 1 vs. 2; female TG, p = 0.026 for day 1 vs. 2). However, percent time freezing did not differ between day 3 (in a modified context) relative to day 1 in female NTG and TG mice. No differences in motor activity were observed within-group across days.

### Unique neuronal and oligodendrocyte gene signatures associated with APP-PS1 familial AD mutations in female mice but not in males

Analysis of DEGs revealed a substantially elevated number of gene signatures in TG relative NTG female mice (Figure 4a). This included 216 upregulated and 24 downregulated genes (log2FC > 1.0, adjusted p < 0.05). DEGs in male mice included 54 upregulated and 5 downregulated genes (log2FC > 1.0, adjusted p < 0.05) (Figure 4b). We used two-way ANOVA to assess cell type composition for normalized gene counts, using geometric means as a composite measure for changes in cell type gene signatures ^64^ (results for Tukey’s post hoc tests are shown in Figure 4c). We observed significant differences in geometric means for microglial (genotype F_1,22_ = 93.5, p <0.0001), endothelial (sex F_1,22_ = 12.78, p = 0.0017) and vascular pericyte (sex F_1,22_ = 5.8, p = 0.02; genotype F_1,22_ = 28.0, p < 0.0001) genes in TG vs NTG mice (Figure 4c, top row). Both male and female mice showed similar geometric means for these gene categories. Conversely, we observed reductions in geometric means for neuronal (genotype x sex F_1,22_ = 8.25, p= 0.0088) and myelinating oligodendrocyte (genotype x sex F_1,22_ = 6.1, p= 0.02; genotype F_1,22_ = 8.9, p= 0.0067) and newly formed oligodendrocyte (genotype x sex F_1,22_ = 5.1, p= 0.03; genotype F_1,22_ = 13.1, p= 0.0015) cell type categories in TG female mice relative to NTG females (Figure 4c, bottom row). A similar non-significant trend was observed for oligodendrocyte precursor category. This was not observed in male mice (Figure 4c, bottom row).

**Figure 4.**
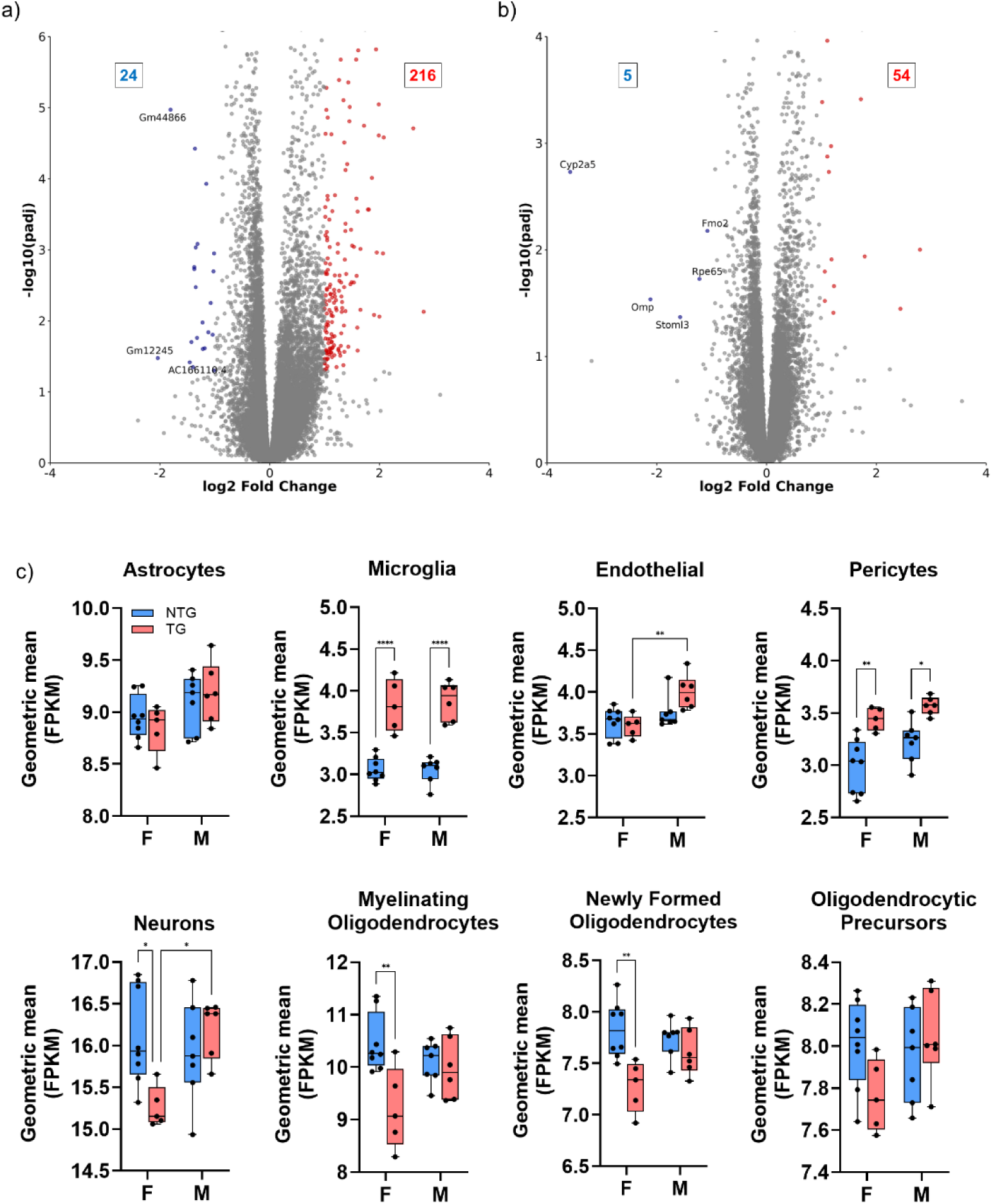
Diverging patterns in differentially expressed genes (DEGs) in female and male APP-PS1 mice. a) Volcano plot showing significant downregulated (blue numbers) and upregulated (red numbers) DEGs as a function of APP-PS1 genotype in female mice. b) Volcano plot of DEGs in male mice. c) Comparison of gene expression signatures for cell type composition groupings between NTG and TG female (F) and male (M) mice. Data are shown as box-whisker with median, minimum-maximum values and individual data points. Asterisks are *p < 0.05, **p < 0.01, *** p < 0.001.

These results highlighted an interesting relationship between amyloid and neuronal and white matter related gene signatures, specific to female mice. WGCNA of RNAseq data revealed 10 modules with significant positively correlated genes and 7 modules with significant negatively correlated genes between TG vs NTG female mice (Figure 5a). Among the identified modules, we further analyzed gene signatures within the brown and lightblue4 modules (Figure 5b-c, respectively). The former is comprised of 669 positively correlated genes (r = 0.91, p = 1.87E-05), of which the top ten included genes related to immune activity (macrophage associated Cd68 was considered the ‘hub’ gene in this module). The latter module is comprised of 439 negatively correlated genes (r = −0.81, p = 6.8E-04), with the top10 most significant genes known for their roles in neuronal and behavioral functions (Gng7 was considered the ‘hub’ gene in this module). Statistical hierarchical cluster heatmaps in Figure 5d-e summarize comparisons within these two modules between TG and NTG female mice. This analysis illustrated a significant segregation of upregulated and downregulated gene clusters between NTG and TG female mice within each module.

**Figure 5.**
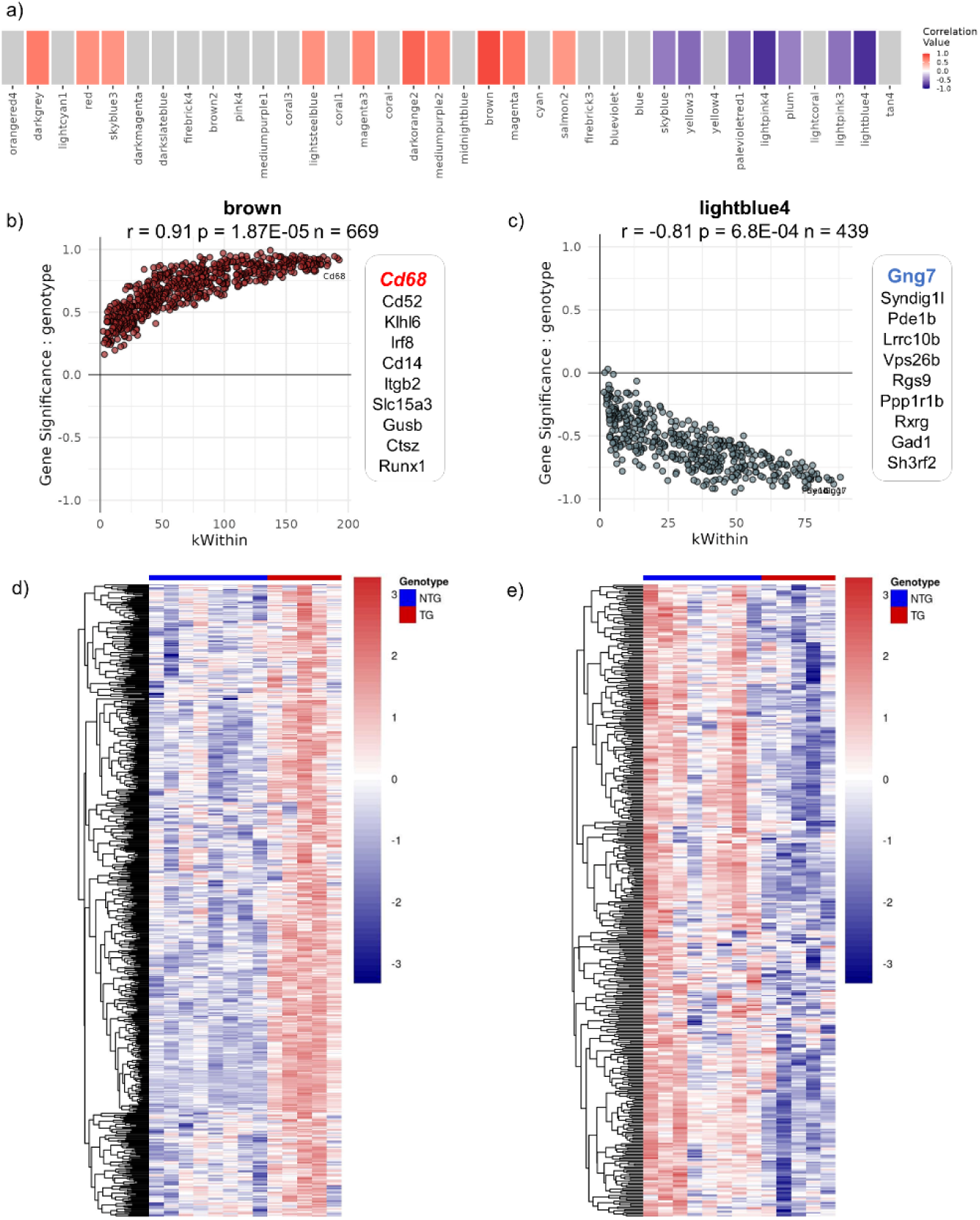
Weighted gene correlation network analysis identified two distinct modules in female APP-PS1 brain with significant positively and negatively correlated groups of immune and neuronal genes. a) Heatmap showing modules with gene networks that are positively (red) and negatively (blue) correlated with genotype (modules with p-value less than 0.1 are indicated by color). Modules with nonsignificant correlations are in grey. b) Gene significance by within module degree (kWithin) for the brown module (evaluated for genotype). Listed are the top 10 most significant genes with hub gene colored red. c) Gene significance by within module degree (kWithin) for the lightblue4 module (evaluated for genotype). Listed are the top 10 most significant genes with hub gene colored blue. d) Hierarchical clustering of gene expression in brown module. e) Hierarchical clustering of gene expression in lightblue4 module.

Module preservation analysis indicated that the brown module containing positively correlated genes was highly preserved between male and female mice (Figure 6a-b). The lightblue4 module containing negatively correlated genes was moderately preserved in male relative to female mice (Figure 6a-b). Consistent with this result, analyses of normalized gene counts for the brown module showed significant increases for the first top 5 genes in both female and male mice (Figure 6c). These included Cd68 (genotype F_1,22_ = 165, p < 0.0001), Cd52 (sex x genotype F_1,22_ = 6.6, p = 0.02), Klhl6 (genotype F_1,22_ = 126, p < 0.0001), lrf8 (genotype F_1,22_ = 93.6, p < 0.0001), and Cd14(genotype F_1,22_ = 27.5, p < 0.0001). Analysis of normalized gene counts for the top ten gene signatures in the lightblue4 module indicated that these were significantly reduced only in APP-PS1 female mice relative to NTG female controls (Figure 6d). These included Gng7 (sex x genotype F_1,22_ = 4.98, p = 0.036), Syndig1l (sex x genotype F_1,22_ = 8.7, p = 0.0075), Pde1b (sex x genotype F_1,22_ = 8.5, p = 0.008), Lrrc10b (sex x genotype F_1,22_ = 7.8, p = 0.01), Vps26b (genotype F_1,22_ = 15.6, p = 0.0007), and Ppp1r1b (sex x genotype F_1,22_ = 9.9, p = 0.004).

**Figure 6.**
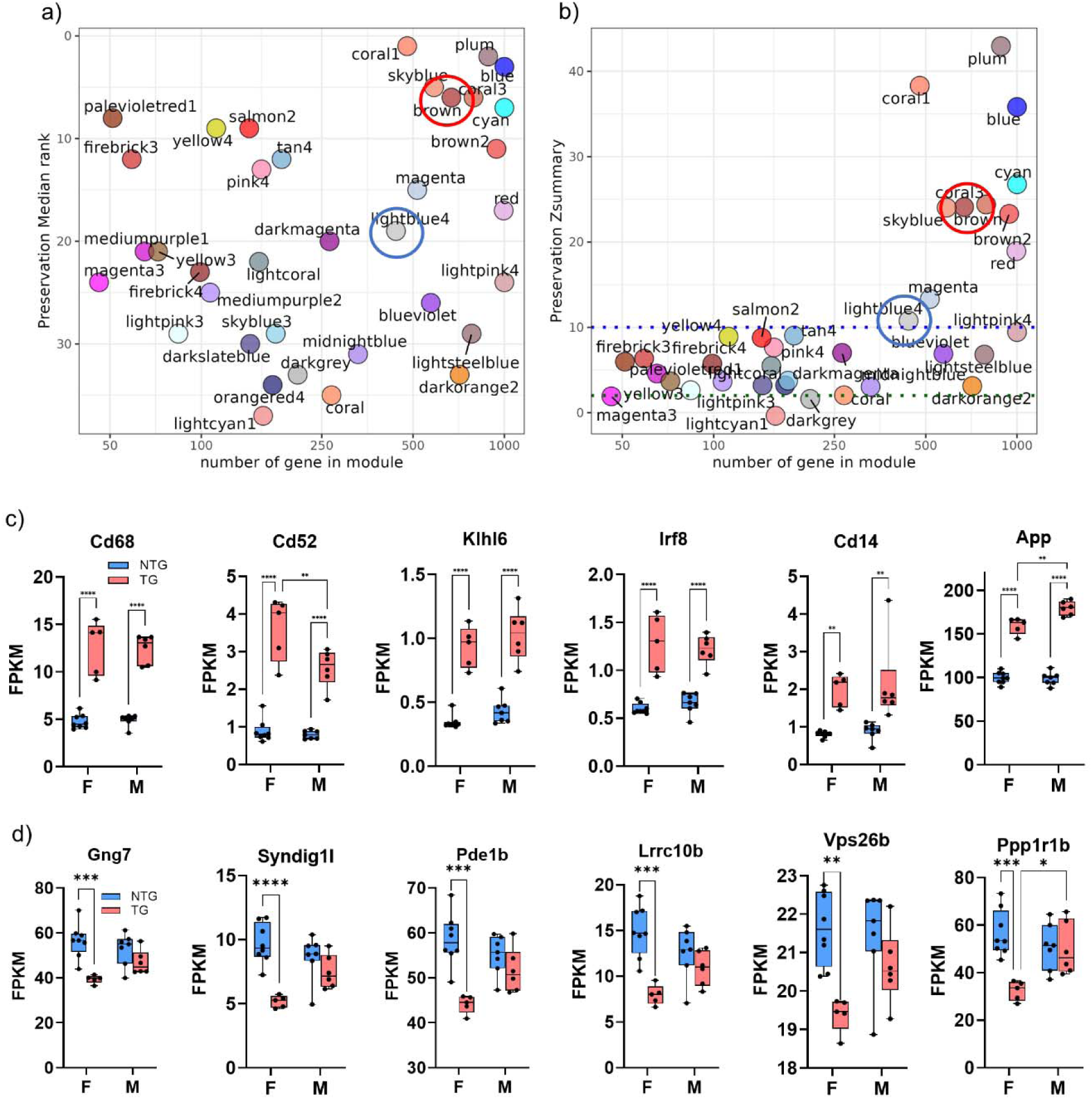
High module preservation of DEGs involved in immune responses (brown module) and moderate preservation of DEGs involved in neuronal function and structure (lightblue4 module). Female module DEGs used as reference to test male DEGs. A) Module rank. Modules close to 0 are highly preserved. B) Zsummary plot. Moderate preservation Z = 2-10 and high preservation Z > 10. d) Comparison of normalized gene expressions for the top 5 most significant genes in the brown module. App gene included as a comparison. e) Comparison of normalized gene expressions for the top 5 most significant genes in the brown module. Ddx3x gene with sex dependent expression levels is included as a comparison. Data are shown as box-whisker with median, minimum-maximum values and individual data points. Asterisks are *p < 0.05, **p < 0.01, *** p < 0.001, ****p < 0.0001.

### Amyloidosis is associated with similar microglia abundance in male and female mice

Our histopathological analysis also confirmed the presence of Iba1-labeled microglial cells, which appeared in greater abundance in TG compared to NTG mice (Supplementary Figure 4a). Iba1 labeling was greater in hippocampus (p = 0.0004), somatomotor (p < 0.0001) and prefrontal cortex (p = 0.0008) in male and female TG mice relative to their corresponding sex matched NTG controls (Supplementary Figure 4b). 2-way ANOVA with main effects and interaction of sex and genotype was used to determine statistical significance.

## Discussion

In this study we examined the effects of AD-amyloidosis on brain wide functional connectivity and gene expression in middle-aged mice. Our data reveal brain network and gene network correlates involved in the complex relationship between Aβ, functional connectivity, and learned fear in the APP-PS1 mouse. These mice develop hippocampal Aβ deposits by 6 months of age and abundant diffuse and dense core plaques throughout the cortex and hippocampus by 9 months of age^32^. Based on substantial evidence of Aβ plaque induced changes in synaptic activity and plasticity in neocortical and hippocampal areas ^7, 8, 10, 11, 65–68^, we hypothesized functional connectivity differences as a result of amyloidosis in the APP-PS1 mouse. Surprisingly, functional connectivity in well-known mouse ICA networks ^61, 63, 69, 70^, including somatosensory, motor, anterior and posterior cingulate cortices known to develop abundant extracellular plaques in this AβPP mouse strain ^71^, did not differ between TG and NTG mice. The lack of effects of amyloid on functional connectivity is likely due to the mechanism of Aβ aggregation in the APP-PS1 mouse. ΔE22Aβ mice with intracellular aggregation of Aβ and PSAβPP mice with parenchymal plaque deposits showed either no differences in functional connectivity or exhibited mild differences compared to control mice ^30^. On the other hand, Arctic AβPP mice (ArcAβ) with vascular and intracellular Aβ aggregates had significantly reduced cortical functional connectivity in mice ^30^. It will be important in future imaging studies to clarify how specific forms of Aβ aggregation affect neuronal circuit activity and large-scale functional connectivity networks.

Despite no observable effect of amyloid on established mouse ICA networks, we did observe differences in functional connectivity between NTG and TG mice within a mixed striatal-amygdala network. This ICA network was distinct in that it did not clearly fit within previously reported networks in mice^28, 62, 63, 72^. Rather it combines regions and structures not thought to be network specific. This is a result of interactions between regions not classically defined in networks which may have been due to sex and transgene specific activity changes. While this network is speculative at this time, ICA decomposes data based on intrinsic connectivity, meaning the statistical separation of this network is not likely to be an artifact of our processing or scan protocol. Peak z-statistic values were observed in voxels located in posterior striatal-basolateral amygdala. The brain areas showing functional connectivity within this network included sensorimotor, cingulate, ventral hippocampal, midbrain and cerebellar regions. The diffuseness of the brain areas included in this ICA component somewhat parallel the similarly diffuse distribution of Aβ plaques in APP-PS1 mice. Within this ICA network, TG mice had higher functional connectivity than NTG mice in motor cortex, orbital cortex, insular cortex, temporal cortex, entorhinal cortex, dorsomedial striatum and midbrain. A shared feature of these brain areas is that they have either first or second order projections to or from the hippocampus, particularly ventral CA1 ^73–77^. A possible mechanism underlying the coactivation of structures in this ICA component, and the observed amyloid related differences in functional connectivity, is the compensatory plasticity of local circuits and projection neurons of ventral hippocampus that may occur in response to extracellular Aβ plaques ^78^. Such compensatory increases in functional connectivity are likely to fail at much later ages since prior evidence indicates that at 18 months of age interhemispheric fMRI connectivity is reduced in APP-PS1 male mice relative to C57BL6/J wildtype mice ^28^. We should note that hypersynchronous activity in parietal and temporal cortical areas was reported in 18-24 months of age 3xTg-AD mice ^79^. This was correlated with tau but not Aβ burden ^79^. This result in 3xTg-AD mice differs from observations of Aβ associated hyperexcitability in fMRI connectivity in AD, which did not correlate with tau ^80^.

DEG analysis suggested that the observed differences in functional connectivity are linked to changes in neuroimmune activity, particularly microglia activation. Analyses of gene enrichment modules revealed gene clusters playing roles in the immune response, cell surface adhesion and receptor-mediated signaling, all elevated in conjunction with elevated human mutant AβPP in male and female mice. This was consistent with histological data showing similarly elevated microglial Iba1 immunostaining in cortical and hippocampal tissue in male and female TG mice. This is an expected result of amyloidosis, which has been reported in AD and across studies using AβPP mouse strains. It is also consistent with timelines of expression reported for *in vivo* PET studies of 12-13 months of age APP-PS1 using radioligand with affinity for the translocator protein (TSPO) ^81^. Interestingly, temporal cortical TSPO but not Iba1 follows a staged increase pattern similar to Aβ in immunolabeling studies of postmortem AD cases ^82^. Whether the elevated abundance in microglial cells contributes to the reorganization of functional connectivity is unclear. Microglial cells are known to play a phagocytic role in synaptic pruning ^83^, particularly of unused or damaged dendritic spines ^84^, and this in turn may contribute to optimization of neural circuit processing ^83, 85–87^. Mice deficient of the Crx3cr1 chemokine receptor have reduced microglia and this is associated with reduced functional coherence between the prefrontal cortex and hippocampus and reduced functional connectivity ^85^. Crx3cr1 knockout mice also exhibit reduced synapses in adult born dentate gyrus cells along with structural changes in neurons within this region ^88^. Iba1 deficiency in the Allograph inflammatory factor1 (Aif1) knockout mouse reduces hippocampal CA1 spines and pyramidal firing frequencies ^89^. Conversely, lifelong depletion of microglia in mice with deletion of the fms-intron regulatory element (Csfr1^ΔFIRE/ΔFIRE^ mice) showed no differences in the number of excitatory and inhibitory synapses nor spine density in the hippocampal CA1, although neurons within this region did show deficits in excitability and altered synaptic properties ^90^.

The observed Aβ related changes in functional connectivity and DEGs were not accompanied by differences in fear conditioning between TG and NTG mice, which was unexpected given previous behavioral characterization of the AβPP mouse ^34, 91, 92^. We did not observe differences between NTG and TG mice on any of the test days, although a trend towards lower freezing responses in male TG mice was observed on day 1. This finding suggests that the observed brain functional connectivity differences precede cued and contextual memory differences, a result that may be specific to the mouse APP-PS1 strain used in this study. It is worth noting that changes in functional connectivity and Aβ plaque related PET signals have been reported in elderly individuals without cognitive deficits ^93, 94^. Although this seems plausible in the present mouse imaging study, there are alternative explanations for the lack of differences in fear conditioning that requires consideration. The combined classical-contextual fear conditioning assay (tone-context-shock pairing) and the moderate-to-high current level for shock delivery used in the present study could account for differences relative to previous work using a standard contextual fear conditioning assay (context-shock pairing) ^34^. As discussed in recent work by our group ^36^, age- and strain-related declines in sensorimotor function can impact performance on fear conditioning assays and the outcomes may reflect underlying mechanisms not directly related to impaired learning and memory. The hybrid classical-contextual fear conditioning assay using auditory plus contextual cues, and a higher current level than in other studies (typically in the range of 0.45-0.6mA over 2 sec), was used here to produce robust training in APP-PS1 mice. This approach may have masked mild cognitive impairment observed with other cued and contextual conditioning protocols that have been used in this strain of AβPP mice. It is worth noting that previous studies using the same APP-PS1 strain tested on a cued/contextual fear conditioning protocol have not observed differences compared to wildtype littermate controls at 8-9mo^95^, although differences are more consistently observed at 13-14 month APP-PS1 mice^96–98^. Conversely, others have reported differences in long-term contextual fear (cited as either 24h or 30 days post-training) between APP-PS1 and controls at 2-6 months of age^33, 92, 99^, with the latter age group showing no short-term differences at 1-hour post-training^92^. Thus, thus the lack of behavioral differences in relation to differential gene expression patterns and functional connectivity remain difficult to explain currently, whereas there seems to be a closer correspondence between functional connectivity, Aβ and DEGs.

Our results reveal significant sex dependent effects of amyloidosis on histopathological, gene expression, functional brain activation and fear conditioning measurements. This is consistent with previously reported sex differences in this mouse model^100^. We should note that while the Prnp promoter has been associated with sex dependent patterns in gene expression in peripheral tissues, this has not been observed for brain^101^. Analysis of the sex dependent gene Eif2s3x^102^ alongside App and Prnp in the present cohort of mice illustrates that the latter genes were not differentially expressed by sex in NTG mice (Figure 6c and Supplemental Figure 5). We observed sex differences in eigenvector centrality, a measure of node ‘hubness’, which was differentially altered by amyloidosis in female compared to male mice. The brain regions with highest eigencentrality in both male and females included the anterior thalamus, dorsal striatum and areas of medial prefrontal cortex and orbital cortex (illustrated in Figure 5c). Functional connectome eigenvector centrality was lower in several prefrontal areas and striatum of NTG female mice relative to NTG males. Sex differences in centrality in functional connectivity networks have been reported in human cortex ^23^ but in no other rodent fMRI studies to date. In humans, women reportedly have higher eigenvector centrality scores in hippocampus and posterior cingulate cortex than men ^23^. We tested another centrality measure, betweenness centrality, which on average resulted in essentially identical distribution of node betweenness centrality rankings across male and female mice (data not shown). This suggests that the observed sex differences in eigencentrality are not generalized to other network centrality measures. It is unclear if this topological organization has a neurophysiological underpinning relevant to cognitive or emotional states in humans and other species.

AβPP female mice had higher eigenvector centrality values in prefrontal cortical regions than controls. Indeed, eigenvector centrality in AβPP female mice were comparable to TG and NTG male mice. This may have significance to female-specific effects of amyloid in modulating the functional and behavioral roles of prefrontal brain regions. This is supported by research showing that PET-based amyloid positivity is related to increased eigenvector centrality in prefrontal cortex, even in the absence of declines in cognitive performance ^22^. The link between amyloid positivity and eigenvector centrality in cognitively healthy controls appears to be relatively specific to the default mode network ^18, 22^. The increased eigenvector centrality observed in posterior and mid-cingulate cortical regions correlates with elevated cerebral spinal fluid (CSF) markers of Aβ and phosphorylated tau (p-tau) ^21^ and this in turn relates to cognitive impairment ^21^. Others have reported differences in eigenvector centrality linked to stable versus progressive mild cognitive impairment ^103^. Eigenvector centrality in the lateral superior temporal gyrus changes with age and may inform on cognitive decline in MCI ^19^. Overall, there is evidence from human neuroimaging studies supporting the observed sex differences in eigenvector centrality and a potentially important link between this network measure and amyloidosis.

In support of the observed effects of amyloidosis on eigenvector centrality in female mice, we found that amyloidosis decreased a cluster of genes specifically in female mice. While the amyloid and immune related genes were highly preserved in modular analyses, the gene clusters reduced in APP-PS1 females was moderately preserved when comparing to male mice. This female specific pattern of gene expression changes included Gng7, Syndig1l, Pde1b, Lrrc10b, Vps26b as the top 5 downregulated genes in AβPP females, and Gad1 (glutamic acid decarboxylase 1 gene encoding the main enzyme involved in producing GABA in neurons) within the top 10 downregulated genes. *Gad1* is also known as Gad67 and is found widely distributed in inhibitory interneuron cell soma ^104^. AD associated alterations in GABAergic neuron function and structure in hippocampal, temporal and parietal regions are well published ^6, 8^. Aβ is linked to hypersynchronous field potential activity in mice via a voltage gated sodium channel – Nav1.1, mechanism in Gad67-expressing parvalbumin interneurons in parietal cortex of mice ^10^. An intriguing notion is that lightblue4 module genes are linked to functional dysregulation or loss of GABAergic neurons in response to Aβ, specifically in female mice. *Gng7* reportedly harbors differentially methylated positions associated with Braak stage pathology ^105^ and in mice has been causally linked to ischemic stroke induced motor dysfunction and stressor-induced anxiety ^106^ and striatal adenosine-A_2A_ receptor-containing medium spiny neurons’ (MSNs) control over locomotor responses to psychostimulants ^107^. Interestingly, *Gng7* is one of the target genes of microRNAs, miR-881-3p and miR-504, that are downregulated by alcohol treatment in female but not male rats ^108^. *Syndig1l* harbors epigenetic CpG-related single nucleotide polymorphisms associated with AD in Hispanic populations ^109^. *Pde1b*, encoding a phosphodiesterase 1, has been linked to motor activity and spatial learning ^110^, identified in motor control in striatum of transgenic mice of Huntington’s ^111^ and involved in depression-like behaviors in response to stress in mice ^112^. Overexpression of *Pde1b* in dopamine D1 receptor-containing striatal MSNs attenuated the locomotor response to cocaine in female mice but not males ^113^. *Ppp1r1c* also known as Darpp32, is found in dopamine receptor D1 expressing striatal medium spiny neurons ^114^. Darpp32 follows a sex-dependent pattern of expression ^115, 116^ and its expression has been shown to be altered by estradiol ^117^. These new data offer useful insight into potential avenues to investigate sex-specific and hormonally regulated mechanisms of amyloid aggregation that may differentially impact female brains compared to males.

Finally, we observed a higher number of extracellular plaques in female versus male mice. This was observed in somatomotor cortex and basolateral amygdala. These findings are consistent with previous research using the APP-PS1 mouse ^118, 119^, the Tg2576 ^120^ and the triple transgenic AD mouse (3xTg-AD) ^121^. The hippocampal concentrations of Aβ40 and Aβ42 and number of plaques at 12 and 17 months of age is more than twice the amount in females compared to male APP-PS1 ^118^. The mechanism leading to higher plaque numbers in female versus male mice may involve effects of estrogen loss on Aβ overproduction. Ovariectomy increases Aβ levels in female Tg2576 mice (measured in whole brain homogenates by mass spectrometry), and estrogen replacement reduces Aβ to sham Tg2576 control levels ^122^. In that study, estrogen did not appear to alter full length human AβPP, secreted AβPP nor the PS1 C-terminal fragment ^122^. A similar increase in Aβ load following OVX, and a reduction with estrogen treatment, is observed in CA1, subiculum and frontal cortex of 3xTg-AD mice ^123^. Estrogen appears to repress BACE1 gene expression, and its activity, via estrogen receptor alpha ^124^, a mechanism that may underlie the higher number of Aβ plaques in AβPP female versus male mice ^122^. Results for gene enrichment analysis further substantiated the observed sex differences in amyloid expression, unexpectedly revealing a diametrically opposed App gene levels compared to the histological data. App gene counts were higher in males compared to female mice, whereas the opposite was found with Aβ plaque immunolabeling. This is consistent with a reduction App gene expression in TG females because of putative estrogenic repression of BACE1 ^124^.

We note several limitations of our studies. First, the use of anesthetics and sedatives for imaging is always a limiting factor. However, the protocol used here, which involves low isoflurane following a continuous infusion of dexmedetomidine, has been well established and used widely across rodent imaging studies. Second, whether the connectome measures correspond to neuronal activity changes versus vascular or astrocytic differences between TG and NTG male and female mice remains unknown and was not addressed here. Finally, while the DEGs were collected in age, strain and sex matched mice, it was collected in a separate cohort of mice from those imaged. Importantly, we did not include protein assessments in this initial study but expect to pursue the role of the identified DEGs in future work.

To conclude, the present study adds to the literature on the functional network changes in response to increased presence of extracellular plaques. Reducing extracellular plaque pathology could be one way in which neuronal communication might be safeguarded in individuals at risk of cognitive decline and Alzheimer disease. Moreover, the identified functional connectome marker, specific to APP-PS1 females and consistent with the human literature, will be interesting to further investigate in other AβPP strains. It is important to consider that the functional connectome measures described here rely on the integrity of neurovascular function. Transgenes under the control of the prion promoter drives protein expression in both neurons and astrocytes^32^, which could potentially affect astrocytic functions in relation to neurovascular coupling in APP-PS1 mice. Furthermore, previously reported Aβ accumulation along the epithelial lining in blood vessel walls^125–128^ is also likely to affect spontaneous fluctuations in BOLD that underpin functional connectivity networks, although whether this occurs in a brain-region-specific manner is unknown. Uncovering how Aβ affects both neuronal networks and vasculature will be critical to narrowing down key contributing mechanisms to functional network changes in aging and AD. Comparing static versus dynamic connectome measures as a function of Aβ expression might be central to this objective^129^. As indicated above, nuanced mechanisms of Aβ aggregation may be important to understand in terms of how this impacts broader network activity.

## Acknowledgements

The authors would like to thank Marjory Pompilus for assistance with portions of this research. Marcelo Febo and Zachary Simon are thankful for the support provided by the McKnight Brain Institute of the University of Florida.

## Author contributions

Zachary Simon (Data curation; Visualization; Writing - Original Draft Preparation; Conceptualization; Methodology; Formal Analysis; Investigation); Karen N. McFarland (Writing - Review & Editing; Conceptualization; Methodology; Formal Analysis); Todd E. Golde (Writing –

Review & Editing; Conceptualization; Methodology); Paramita Chakrabarty (Writing - Review & Editing; Conceptualization; Methodology; Formal Analysis); Marcelo Febo (Data curation; Visualization; Writing – Original Draft Preparation; Conceptualization; Methodology; Formal Analysis).

## Statements and declarations Ethical considerations

All procedures received prior approval from the Institutional Animal Care and Use Committee of the University of Florida and follow all applicable NIH guidelines.

## Consent for publication

All authors approve of the final version of this manuscript.

## Data availability

The TAUAPPms RNAseq datasets used in this manuscript has been previously deposited in AD Knowledge Portal (https://adknowledgeportal.org) with the support of NIH/NIA (U01 AG046139 to T.E. Golde, N.D. Price, and N. Ertekin-Taner). The data can be accessed at https://www.synapse.org/ (Synapse ID:syn2580853). The AD Knowledge Portal is a platform for accessing data, analyses, and tools generated by the Accelerating Medicines Partnership (AMP-AD) Target Discovery Program and other National Institute on Aging (NIA)-supported programs to enable open-science practices and accelerate translational learning. All MRI datasets are uploaded to openneuro.org.

