## Supplemental Materials for "Sex dependent effect of amyloidosis on functional network ‘hub’ topology is associated with downregulated neuronal gene signatures in the APP*swe*/PSEN1dE9 double transgenic mouse" (from another version of this preprint: version 3, 2025-05-22)

**Supplementary figures and figure captions**

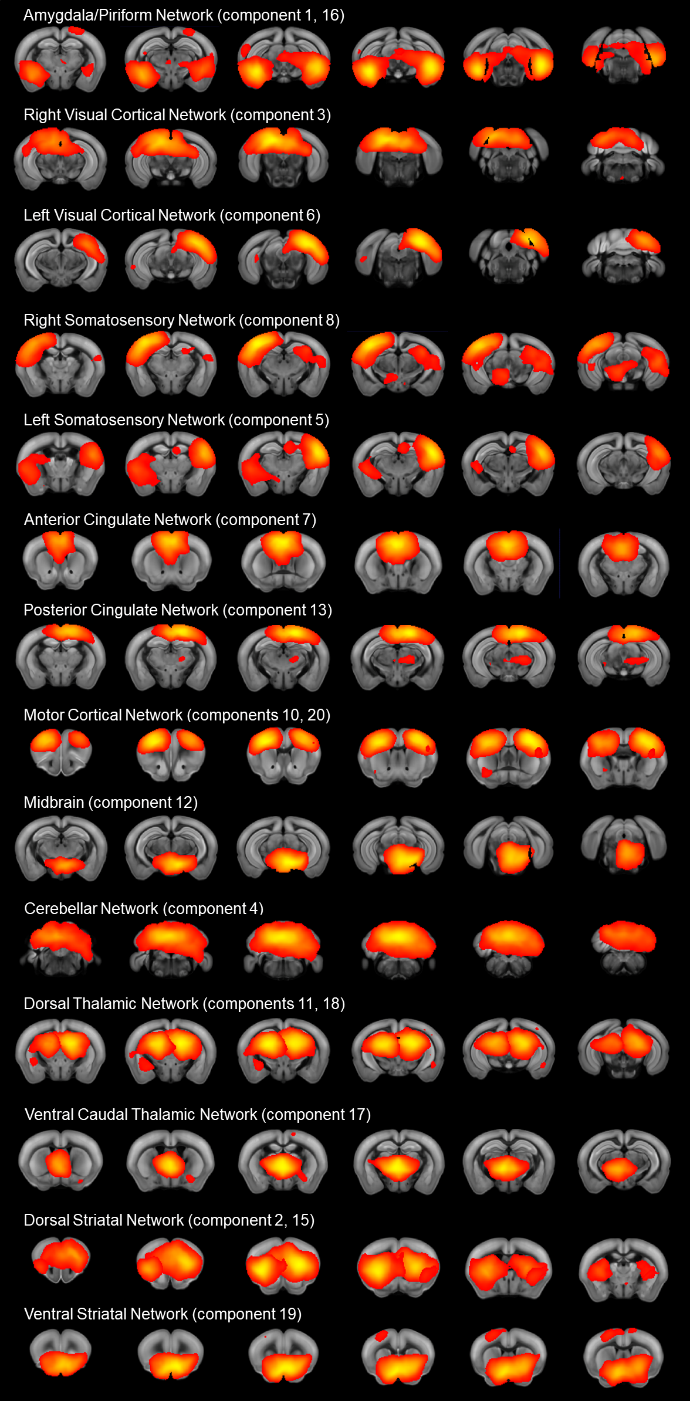

**Supplementary Figure 1.** Independent component analysis of 28 mouse fMRI scans yielded 20 distinct networks. Networks were identified based on prior studies and on location of peak z statistic voxels (higher intensity/higher functional connectivity voxels highlighted in yellow and lower intensity in orange hue). Assigned network names are above each row of ICA maps, with component numbers in parenthesis. ICA maps are organized rostral to caudal from left to right. Z statistical maps are thresholded using threshold free cluster enhanced approach and are overlaid on to the Allen Mouse Brain Common Coordinate Reference atlas (CCFv3).

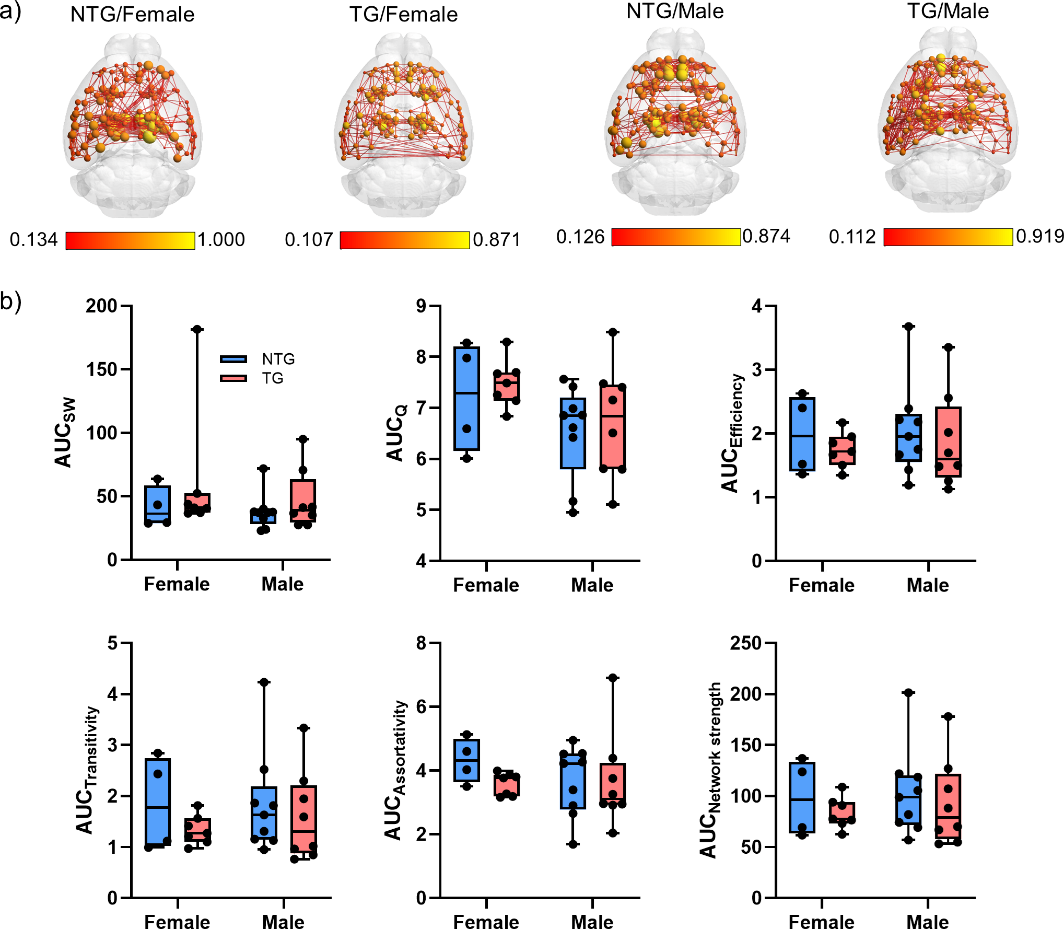

**Supplementary Figure 2.** No significant differences in globally estimated graph theory measures. a) Functional connectome maps averaged within groups and illustrate node strength and edge strength (Pearson correlations) as spherical nodes and lines connecting nodes. Node strength is reflected in sphere size and color intensity (scale bar below maps indicate Fisher z-transformed r values rescaled between 0 and 1). Edges are thresholded at 6% density for visual purposes. b) Comparison of several network measures (converted to area under the curve for network densities between 2-40%) between male and female NTG and TG mice.

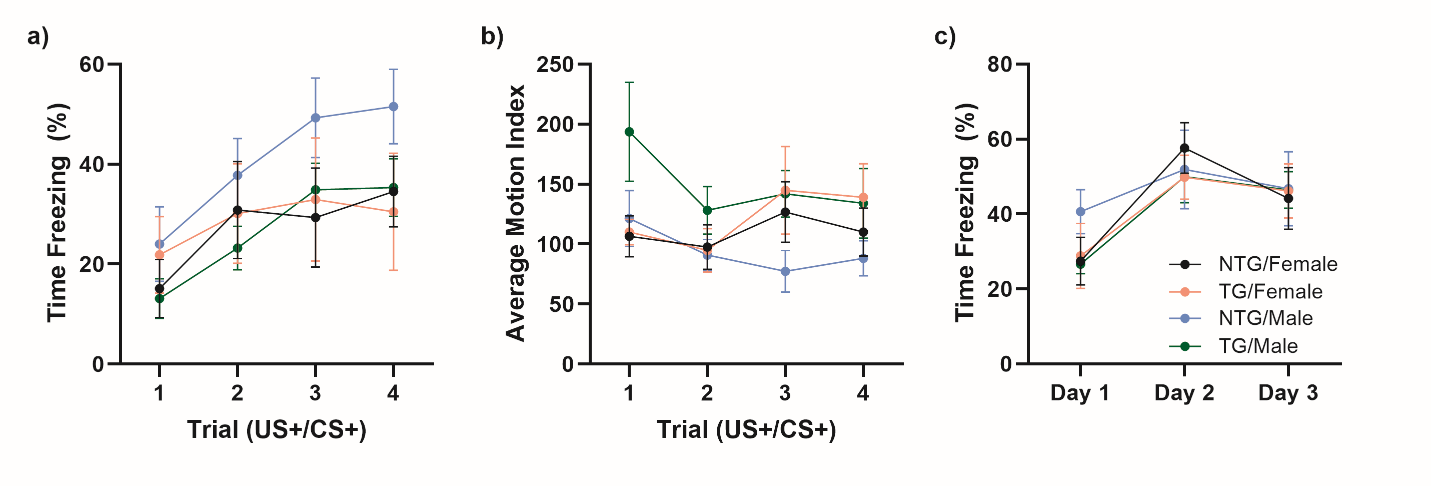

**Supplementary Figure 3.** No significant differences between NTG and TG mice in fear conditioning tests. a) Percent freezing responses per 60 second trials on day one of conditioned stimulus auditory cue/context (conditioned stimulus, CS) pairing with a foot shock (unconditioned stimulus, US). b) Motor activity averaged over each conditioning trial on day 1. c) Percent freezing responses averaged over 60 min trial epochs per day. Day 1 represents fear learning, day 2 recall, and day 3 recall in a modified context (on days 2 and 3 auditory cue presented in the absence of shock). Data in plots are presented as mean ± standard error.

**Supplementary Figure 4.** Amyloidosis is associate with an increase microglial abundance in both male and female mice. a) Representative histological Iba1 antibody-stained sections from NTG and TG mouse basolateral amygdala (BLA), prefrontal cortex (PFC), somatomotor cortex (SMC), and hippocampus (HPC). b) Comparison of microglial abundance between male (M) and female (F) NTG and TG mice. Data in a-b are box-whisker showing median, minimum-maximum values and individual data points. Asterisks are *p < 0.05, **p < 0.01.

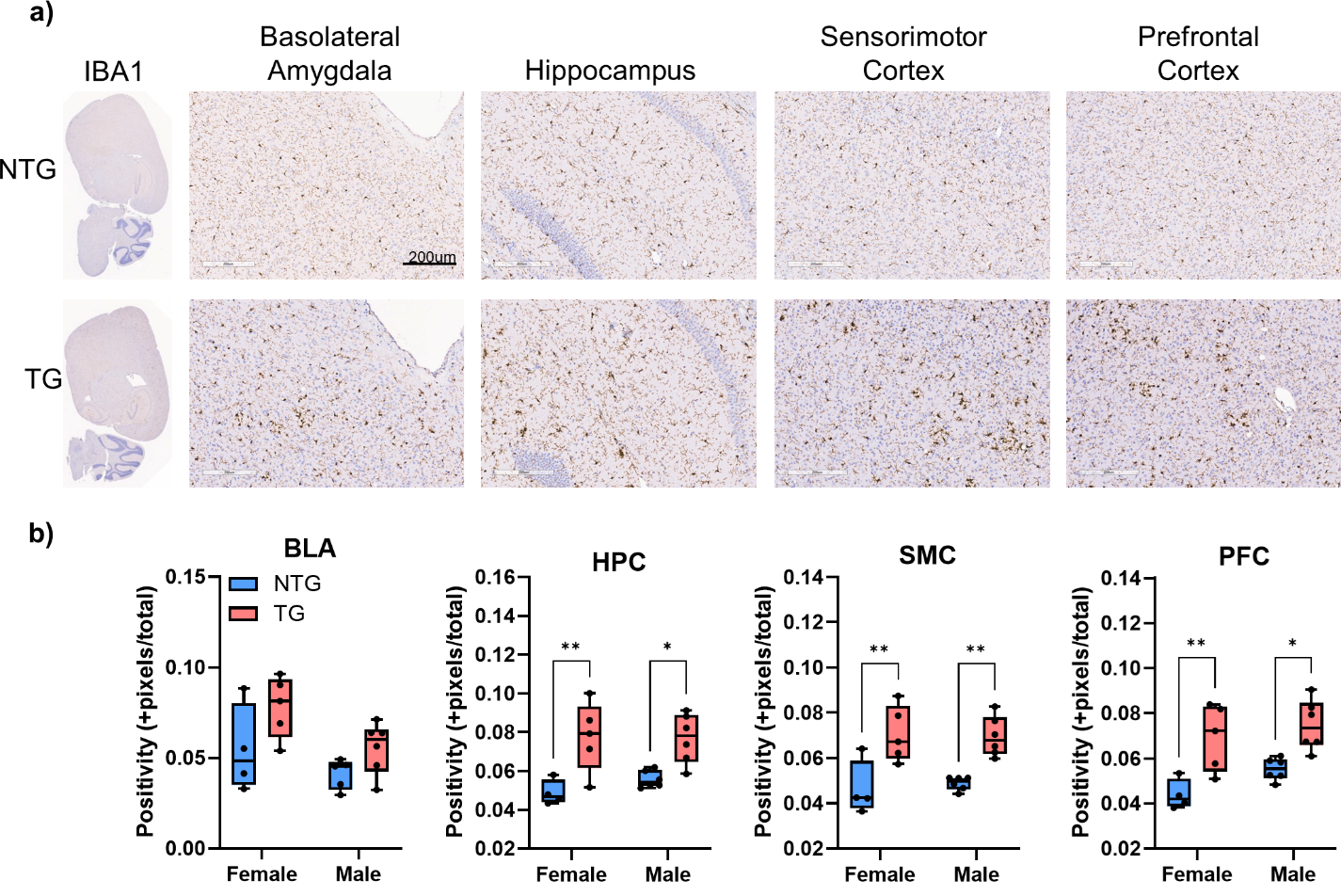

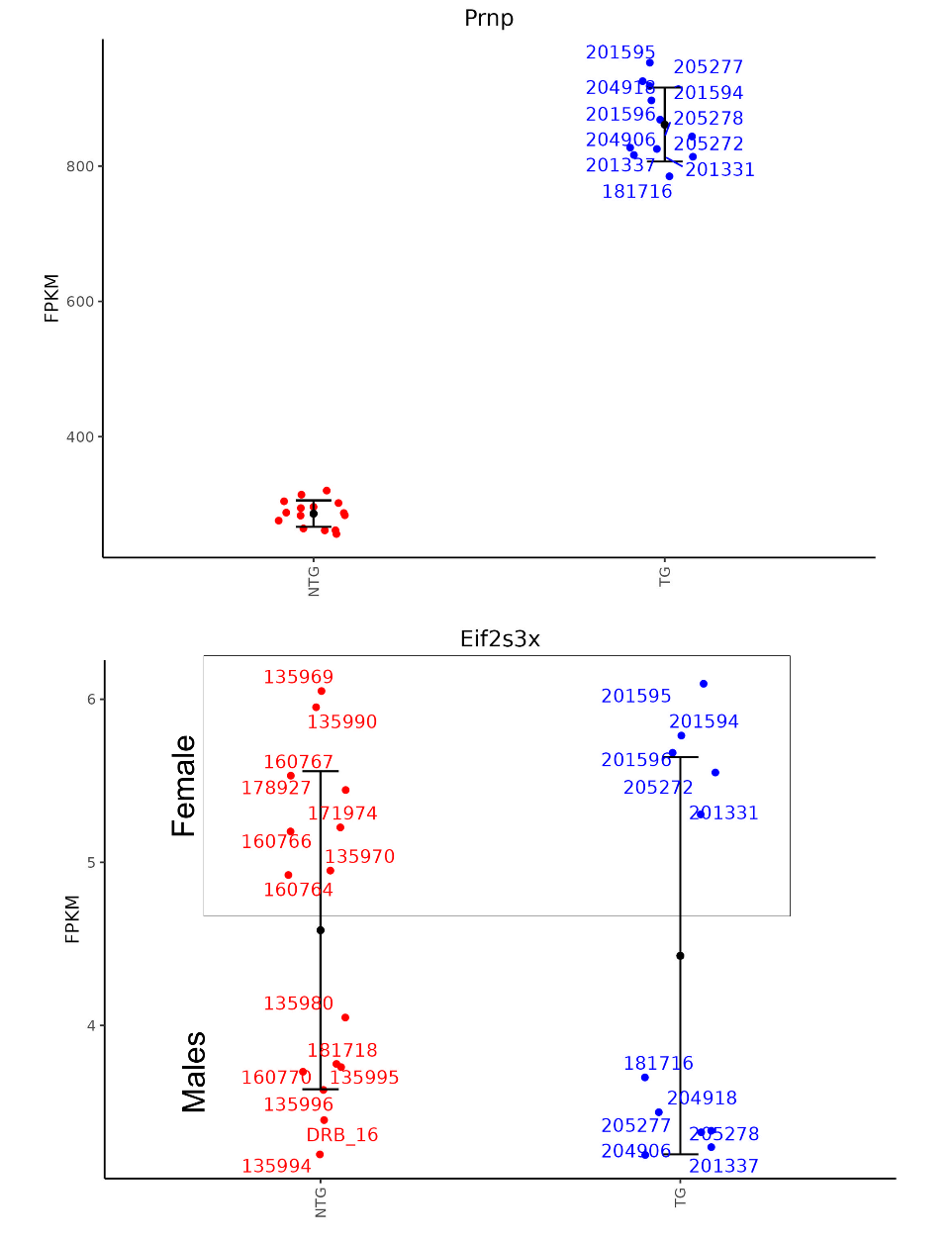

**Supplementary Figure 5.** Comparison of normalized gene expressions for the Prnp gene and a sexually dimorphic gene Eif2s3x. Individual data points are shown and identified according to their subject identifiers (red color dots are NTG and blue TG).
